# Existence and Stability Criteria for Global Synchrony and for Synchrony in two Alternating Clusters of Pulse-Coupled Oscillators Updated to Include Conduction Delays

**DOI:** 10.1101/2024.01.11.575222

**Authors:** Ananth Vedururu Srinivas, Carmen C. Canavier

**Author notes:** **Corresponding Author:** Carmen Canavier.

## Abstract

Phase Response Curves (PRCs) have been useful in determining and analyzing various phase-locking modes in networks of oscillators under pulse-coupling assumptions, as reviewed in Mathematical Biosciences, 226:77-96, 2010. Here, we update that review to include progress since 2010 on pulse coupled oscillators with conduction delays. We then present original results that extend the derivation of the criteria for stability of global synchrony in networks of pulse-coupled oscillators to include conduction delays. We also incorporate conduction delays to extend previous studies that showed how an alternating firing pattern between two synchronized clusters could enforce within cluster synchrony, even for clusters unable to synchronize themselves in isolation. To obtain these results, we used self-connected neurons to represent clusters. These results greatly extend the applicability of the stability analyses to networks of pulse-coupled oscillators since conduction delays are ubiquitous and strongly impact the stability of synchrony. Although these analyses only strictly apply to identical oscillators with identical connections to other oscillators, the principles are general and suggest how to promote or impede synchrony in physiological networks of neurons, for example. Heterogeneity can be interpreted as a form of frozen noise, and approximate synchrony can be sustained despite heterogeneity. The pulse-coupled oscillator model can not only be used to describe biological neuronal networks but also cardiac pacemakers, lasers, fireflies, artificial neural networks, social self-organization, and wireless sensor networks.

**AMS Subject Classification:** 37N25, 39A06, 39A30, 92B25, 92C20

## 1. Introduction

Mathematical methods that are commonly utilized for studying synchronization between neural oscillators [1] include weak coupling [2,3], firing time maps [4–7], fast threshold modulation [8,9] and mean field methods, which often use leaky integrate-and-fire neurons [10]. The first three methods can only apply to repetitively firing neurons where individual neurons act as limit cycle oscillators with a baseline oscillation period that can be measured, whereas the fourth method is most useful in the case in which individual neurons are not oscillators but instead emit spikes randomly. The first two methods employ different variants of phase response curves (PRCs). A phase response curve measures the change in length of the oscillation period caused by an input received during an oscillatory cycle. Weak coupling assumes that each oscillator remains very near its unperturbed trajectory which is the limit cycle [11], and the effects of multiple inputs are summed to get the averaged effect over one (unperturbed) oscillation cycle. In weak coupling, the infinitesimal phase response curve is obtained in the limit as the strength and duration of the synaptic input current both go to zero.

Here, we study neural oscillators using firing time maps, under the assumption that coupling between neurons is pulsatile, meaning that the effect of the coupling is brief compared to the period of the oscillators. The perturbation is not required to be small, but the oscillator is assumed to return to its unperturbed, free-running trajectory (back to its limit cycle) prior to receiving any additional inputs. We use neural oscillators for illustrative purposes because neurons emit action potentials that trigger a post-synaptic conductance in the neurons to which they are connected, and the duration of the conductance change is often brief compared to the period of the network oscillations. In contrast to infinitesimal PRCs, phase response curves used for the firing time maps are analogous to spike time response curves [12,13]; the perturbation used to measure the PRC is a post-synaptic conductance waveform, often assumed to be biexponential. Neural oscillators are nonlinear; therefore, the effect of a perturbation is not time invariant, but rather periodic with the same period as the neural oscillator. The PRC is the normalized change in cycle length plotted as a function of the phase of the oscillator. In a network in which each neuron receives N synaptic inputs, any given neuron can receive up to N simultaneous inputs. Since multiple inputs are not assumed to be additive, the PRC to *n* out of N inputs must be measured for each *n* ∈[0, *N*], and cannot be inferred from the single neuron PRC.

## 2. Methods

### 2.1 Conductance-based Model Neurons

To test the methods developed herein on neural oscillators, we used previously published models of inhibitory [14] and excitatory neurons [15]. Detailed description of these models can be found in Appendices A1 and A2 respectively. In order to make the inputs more pulsatile and reduce deviations from the assumptions given below in section 2.4, we reduced the decay time constant of the biexponential inhibitory synapses from (*τ*_1_) 2.0 ms in [14] to 0.45ms for illustrative purposes.

### 2.2 Phase response curves

We use a variant of the phase response curve (PRC) also called a spike time response curve [16] because the perturbation used to generate the PRC is the conductance waveform in the target neuron elicited by a spike in its presynaptic partner. These PRCs are measured using the protocol described in Figure 1A. The inset at lower left illustrates the open loop condition in which an input from one oscillator is applied at different points in the cycle of another oscillator in the absence of any feedback. The first and second [17] order phase responses *f*_*j,A,B*_*(φ)* are defined as the normalized difference between the perturbed cycle period T_j_ and the free-running intrinsic cycle period P_i_ when an input is applied at a phase φ: *f*_*j,A,B*_*(φ)=(T*_*j*_*-P*_*i*_*)/P*_*i*_, where j = 1 or 2. Fig. 1B shows the first and second order PRCs generated using the response of Via model [14] for fast spiking neurons (described in Appendix A1) to a biexponential inhibitory conductance waveform (red trace in Figure 1A). The second and third subscripts on the phase response curves are omitted for simplicity where they are clear from the context. Figure 1A defines the stimulus and response intervals based on the open loop PRC as *ts = P*_*i*_*φ* and *tr= P*_*i*_*(1-φ+f*_*1*_*(φ))*, respectively. The stimulus interval is the time elapsed between the most recent spike and the start of the input waveform. The response interval is the time elapsed between the start of the input waveform and the next spike time. This interval is simply the part of the free-running period remaining when the input is applied plus the phase response.

**Fig. 1.**
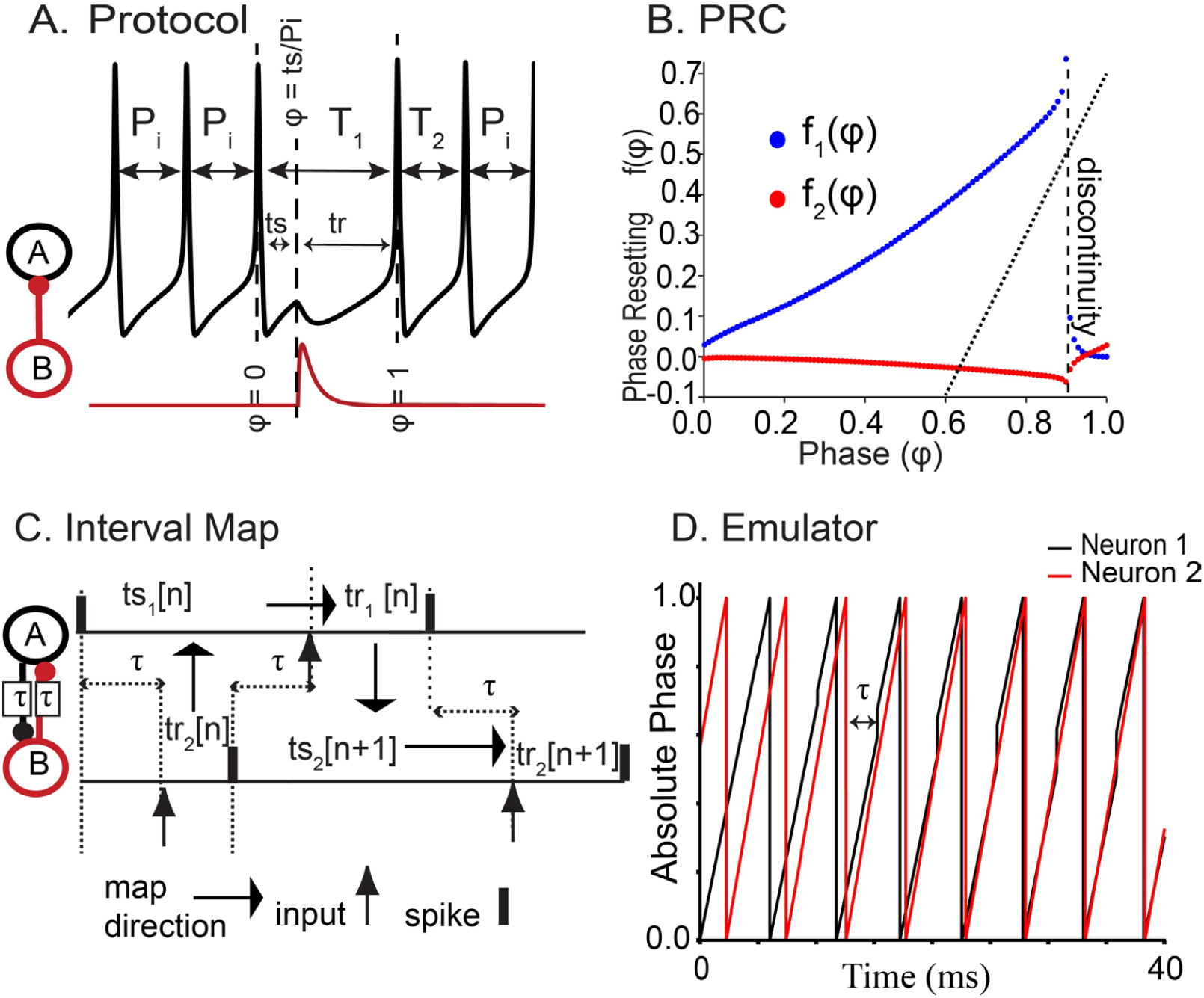
Phase Response Curve Under Pulsatile Coupling Assumptions. A. A single output waveform from one oscillator (red) is used to perturb the other at phases ranging from 0 (spike onset) to 1 (onset of the next spike). The phase φ is the stimulus interval (ts) normalized by the unperturbed period P_i_. B. The phase resetting f_j_(φ) is the normalized change in the cycle length caused by the perturbation (T_j_-P_i_)/P_i_ where j=1 or 2 for first and second order resetting respectively. The response interval tr is determined by the phase resetting tr= P_i_(1-φ+f_1_(φ)). The vertical dashed line indicates a discontinuity and the dotted line is explained in section 3.4. C. The PRC defines an interval map in a two-neuron network with delays. The response interval in one neuron determines the stimulus interval in the other, which determines the next response interval in the same neuron. D. Emulator output (map with no pre-determined firing order). Two neurons converge towards synchrony. The phase of each neuron increases linearly except when reaches its threshold value φ = 1 or when its phase is reset after receiving a delayed input from the other neuron.

### 2.3 Stability Proofs

The basic idea is to assume a firing pattern (for example, the alternating firing pattern with delays in Fig. 1C) and create a discrete system of difference equations (a map or coupled set of maps) that characterize the dynamics in response to a perturbation. Then we linearize the map about the fixed point of the map and find the eigenvalues of the Jacobian matrix. These eigenvalues must have an absolute value less than 1 for stability. The maps rely upon the definition of stimulus and response intervals adapted from the open loop condition in Figure 1A to the closed loop condition in Figure 1C. In the network, the stimulus interval *ts*_*i*_*[n]* in cycle *n* is the interval between when neuron *i* fires and when it next receives an input: *ts*_*i*_[*n*] = *P*_*i*_*φ*_*i*_[*n*], where *P*_*i*_ is the intrinsic period of neuron 1 and *φ*_*i*_*[n]* is the phase at which an input is received on cycle *n*. If two inputs are received in the same cycle, the resetting due to the first input is assumed to be complete by the time the second input is received and the second stimulus interval is *ts*2_*i*_ [*n*] = *P*_*i*_ (*φ* _*j*_ [*n*] −*φ*_*i*_ [*n*] + *f* (*φ*_*i*_ [*n*])), where *φ*_*j*_ is the phase at which the second input is received, and *f(φ*_*i*_) is the phase resetting due to the first input. The response interval *tr*_*i*_*[n]* is the interval between the receipt of the last input and the next spike in neuron *i*: *tr*_*i*_ [*n*] = *P*_*i*_ (1− *φ*_*l*_ [*n*] + *f* (*φ*_*l*_ [*n*])), where *φ*_*l*_ is *φ*_*i*_ if there has only been one input in the cycle and *φ*_*j*_ if there have been two.

### 2.4 Assumptions of pulsatile coupling

Firing time maps assume pulsatile coupling and are based on the phase response curve (PRC) measured under open loop conditions (Figure 1A), meaning that there is no feedback to the rest of the circuit during the measurement. For the PRC measured under those conditions to apply to a closed loop circuit (Figure 1C) with feedback from other circuit elements, several assumptions must be honored: 1) The trajectory of each oscillator must be near its unperturbed limit cycle when an input is received. 2) If there are multiple inputs within a network period, then the trajectory should be near the limit cycle at the time that each input is received. It is not required that the trajectory be near the limit cycle when a spike is emitted. 3) Second order and higher resetting should be negligible. 4) There should be no non-negligible cumulative effect of multiple inputs such as firing rate adaptation [18] or acceleration [19].

### 2.5 *Emulator* (Map with no predetermined firing order [5,20])

We use a hybrid oscillator [21] that updates its phase continuously and linearly between events but discretely when an input is received or its threshold is crossed:

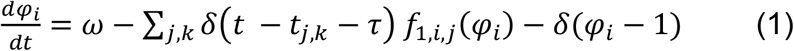

where *φ*_*i*_ is the phase of the *i*^*th*^ neuron, ω is the angular frequency, δ() is the Dirac delta function, *t*_*j,k*_ is the time the *k*^*th*^ input is received from neuron *j, τ* is the synaptic delay and *f*_*1,i,j*_(*φ*_*i*_) is the first order phase response of neuron i at time *t*_*j,k*_ to an input from neuron *j*. The phase is reset to zero when the phase reaches 1 and a pulse is emitted. Since PRCs are not additive, at the time two neurons synchronize their input to neuron *i*, for strict accuracy the appropriate PRC corresponding to the joint input must be substituted for the two separate PRCs. We utilize the emulator to test our proofs because the pulsatile coupling assumptions are honored exactly in the emulator, whereas in simulations using conductance-based models we cannot ensure that the assumption of pulsatile coupling is strictly honored. Figure 1D shows two coupled oscillators characterized by their PRCs converging to synchrony, using the emulator. The input given to this map is the intrinsic period and phase resetting curve of each neuron.

## 3. Previous Work on the Application of Firing Time Maps to Synchronization

We reviewed the application of firing time maps to synchronization of pulse-coupled oscillators comprehensively in 2010 [5]. We summarize the most relevant results here, and also update the summary to include relevant new results [20,22–25] obtained after our previous review [5] was published.

### 3.1 Forced Oscillator

The simplest case of phase locking in pulse-coupled oscillators is that of a single oscillator forcing another oscillator in a one-to-one locking in which the driven oscillator receives an input on each cycle at a phase of *φ*. The stability criterion derived by [26] is |1 – *f*’(*φ*)| < 1, where *f’(φ*) is the slope of the PRC at the locking phase *φ*. The perturbation away from the locking is multiplied by the factor *1 – f’(φ)* on each cycle. For stability, the perturbation must decay to zero; therefore, for stability the absolute value of the factor must be less than one. The slope *f’(φ)* is the ratio of the compensation by the PRC back towards a steady locking to the perturbation from a steady locking. If the *f’(φ)*=1, the factor multiplying the perturbation is 0, and 1:1 locking is restored within a single cycle.

### 3.2 Self-connected oscillator with delay

Previously, a self-connected oscillator has been taken as the equivalent to a network of identical, synchronized, mutually connected neurons [27], hence synchronization tendencies of a self-connector oscillator are of interest. One study derived existence and stability criteria for a periodically spiking neuron forcing itself with a feedback input at a fixed delay after each spike [28]. Considering delays longer than the network period (P_N_) and neglecting second order resetting, the criterion for existence of a 1:1 phase locked mode is *τ/P*_*i*_ = *φ\** + *k+ kf’(φ)*, where *P*_*i*_ is the intrinsic period of the oscillator, or equivalently P_N_=P_i_+*f’(*τ*-kP*_*N*_) which lends itself to a graphical solution for P_N_, the network period, and gives the locking phase as *φ*= (*τ*-kP*_*N*_) [22,23,28]. The criterion for *s*tability [5,22,28,29] is -1/k< f’(*φ*)<1, where the integer *k* denotes the number of network periods wholly contained in the delay. For *k*=0, all lockings are stable because of the self-correcting effect of the fixed delay between a spike and the receipt of an input without any other intervening events.

There are other possible solutions for self-locked oscillator besides the 1:1 locking considered above. At f’(*φ*)=1, the eigenvalues for the firing map of the self-locked oscillator with delays cross the unit circle [22,29], and a period two [30–32] solution which alternates between two interspike intervals of different lengths emerges. Interestingly, as the delays become longer, an increasing number of multistable solutions with different sequences containing the same two intervals arise, termed multistable jittering by the authors [22,23]. This multistability requires a region of the PRC in which f’(*φ*)>1.

### 3.3 Synchrony Between Two Coupled Oscillators with and without delays

The criterion for synchrony between two identical, reciprocally pulse-coupled oscillators without delays depends on the slope of the PRC at the phases of 0 and 1. Phases of 0 and 1 correspond to action potential initiation, but the pulse is emitted only at a phase of 1. In a network with no delays, both neurons receive an input from the other at a phase of 0 (equivalent to a phase of 1). The definition of a cycle length as being between two action potentials (threshold events) may result in a discontinuous PRC. For example, an input applied just before an action potential may delay that action potential, but an input applied after the action potential has already been initiated cannot delay it. Therefore, the left and right slopes at the phase at which an input is received in the case with no delays can be different. The criterion for stability is [33] |(1 – *f’(0*^*+*^)) (1 – *f’(1*^-^))| < 1. If inputs are received in different intervals, then they are multiplicative. However, if multiple inputs are received in the interval between a spike in one neuron and a spike in another neuron, then the compensations are additive; for a two-neuron network the inputs are received by different neurons. Therefore with small delays (less than a network period), the criterion for stability of synchrony in reciprocally coupled identical neurons [24] becomes |*1 – f’(*τ*/P*_*i*_*) – f’(*τ*/P*_*i*_)| *< 1*, or equivalently for two identical oscillators |*1 – 2 f’(*τ*/P*_*i*_)| *< 1*, where *τ* is the delay and *P*_*i*_ is the intrinsic period of the free running oscillator.

### 3.4 Alternating firing pattern in two neuron networks

The existence criteria [24] that determine the steady locked phases at which each cluster receives an input from the other in an alternating firing pattern is *tr*_*A*_[∞*]+2*τ*=ts*_*B*_[∞] and *tr*_*B*_[∞*]+2*τ*=ts*_*A*_[∞] (Fig. 1C), where the stimulus and response intervals in the steady 1:1 phase locked mode are again defined by the phase response curves. If the neurons are identical, the criterion for an exact antiphase mode reduces to *tr*_*A*_[∞*]+2*τ*=ts*_*A*_[∞], which can be simplified to *f(φ*)=2φ* - 1 –2τ/P*_*i*_ [34], considering only first order resetting. This criterion forms a dotted line in the plane of Figure 1B; the antiphase mode only exists if the line intersects the first order phase response curve.

The general criterion for stability of an alternating firing pattern between two neurons [17,35] A and B is |*(1 – f’*_*A,B*_*(φ\**_*A*_*)) (1 – f’*_*B,A*_*(φ\**_*B*_))| *< 1* where φ*_A_ is the phase at which neuron A receives an input from neuron B and φ_B_ is the phase at which neuron B receives an input from neuron A in a one-to-one phase-locked mode. The stability criterion is unchanged for short delays [24].

## 4. Novel Results

### 4.1 Extension of the results for synchrony in an N neuron network without delay to include delays

For a network of neurons each receiving N-1 identical pulse-coupled inputs with no delays, the criteria for stability of synchrony were previously obtained as an extension of the criterion in section 3.3 for two neurons. The idea was to conceptualize the N neuron network as two oscillators, one representing a single neuron and the other representing the remaining N-1 neurons with N-2 self-connections representing the inputs from the other neurons in the presumed synchronous cluster. Thus, the criteria that determine whether synchrony will be re-established after a perturbation of a single neuron from a synchronous cluster in a network with no delays are 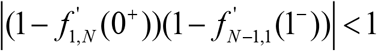 and 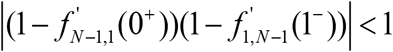 [33]. As before, the indices *i,j* on the phase resetting curve represent the number of neurons in the postsynaptic and presynaptic clusters respectively. Figure 2A shows the reduction of an N-neuron network to a closed loop of two oscillators (with delays added) on the left, and the firing pattern when one neuron is perturbed from the synchronous cluster on the right, including delays. For the two-neuron case, adding even a slight delay changed the criteria for synchrony as given in Section 3.3 [24]. For the N-neuron case, here we find that the delays also change the stability criteria from the case with no delay because both the free-running period and the PRC of the self-connected oscillator depend upon the delay. Figure 2B shows the open loop configurations used to measure the PRCs for the response of each oscillator to an input from the other oscillator. We do not include autapses (self-connections) but the results that we prove below can be easily extended to include autapses. Connections need not be all to all as long as each neuron receives the same number of identical inputs [14].

**Fig. 2.**
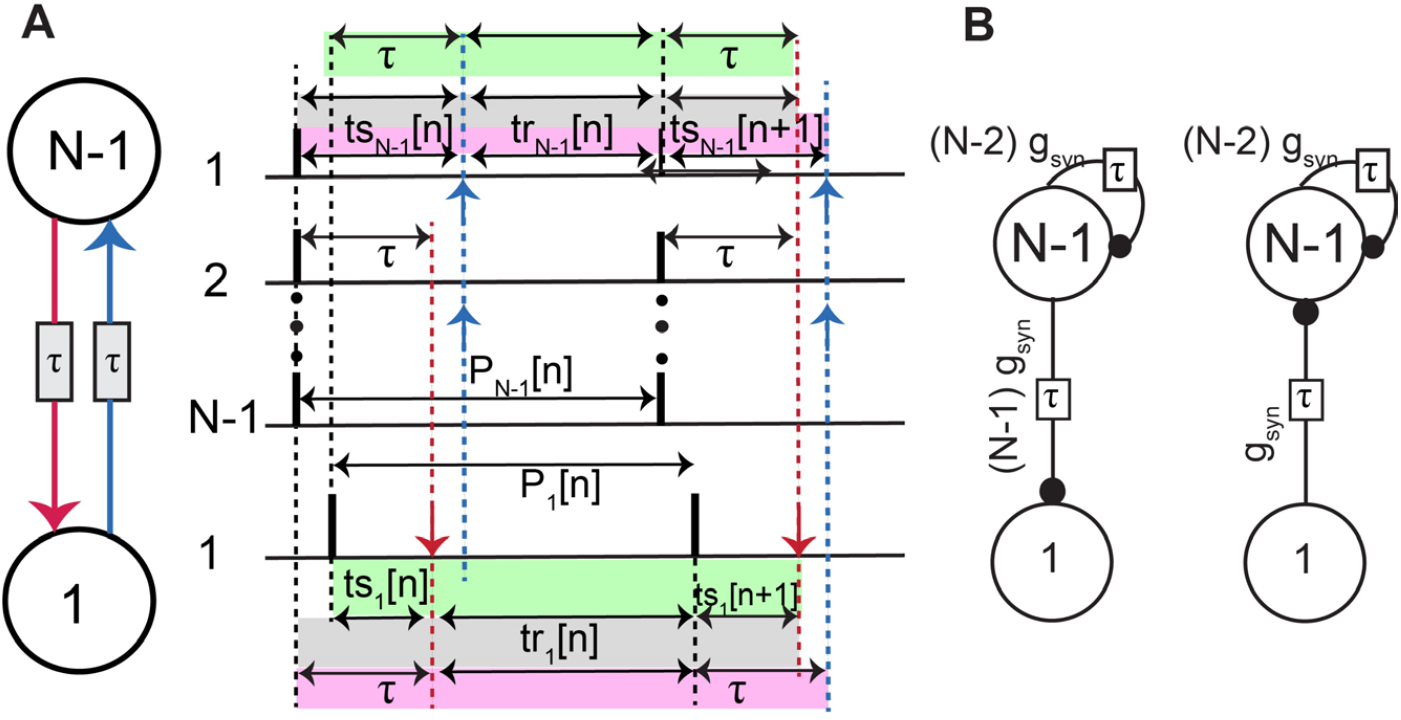
Stability of Synchrony in a Network of N identical neurons each receiving N-1 identical inputs. A. Closed loop circuit (left). Schematic firing pattern in which a single neuron is perturbed from synchrony (right). These intervals are used in the proof of stability. B. Open loop PRCs schematic for reduction of the network to two oscillators. The larger group is assumed to receive delayed synchronous feedback from itself.

For synchrony to be stable in an N-neuron network, if a single neuron is perturbed, the two resulting oscillators (denoted 1 and N-1) must re-synchronize as a single network of N oscillators. To obtain a criterion that describes the stability of synchrony, we first assume a synchronous firing pattern, then assume a single neuron is perturbed from that pattern as in Fig. 2A. The two sets of intervals shaded in each color (green, gray and pink) have equal length resulting in three equations:

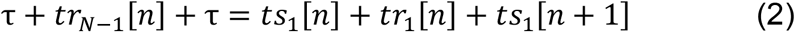

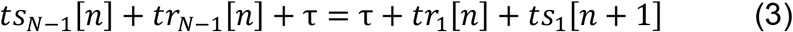

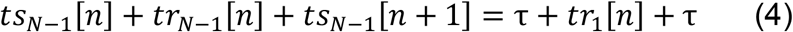

Stability is then determined by the eigenvalue(s) of the discrete map. The discrete map expresses the perturbation of the phase variables of oscillators from the steady state in the [*n*+1]^*th*^ cycle in terms of the same variables in the [*n*]^*th*^ cycle.

Let *P*_1_ be the intrinsic period of the single perturbed neuron and let *P*_*N*−1_ denote the network period of the N-1 unperturbed neurons assumed to be oscillating synchronously. The stimulus and response intervals of the oscillator *i* in its *n*^*th*^ cycle are defined as *ts*_*i*_[*n*] = *P*_*i*_*φ*_*i*_[*n*] and *tr*_*i*_[*n*] = *P*_*i*_(1 − *φ*_*i*_[*n*] − *f*_1,*i,j*_ (*φ*_*i*_[*n*]), respectively where *f*_1,*i,j*_(*φ*_*i*_[*n*]) is the first order PRC of the oscillator *i* due to the input from oscillator *j* in its *n*^*th*^ cycle. Here, *i* and *j* take values 1 and N-1 in this case. We substitute these definitions for the intervals as shown in Appendix B. Since we only use first order PRC in this derivation, we will henceforth denote the PRC functions as *f*_*i,j*_ for convenience.

We define the phase in the n^th^ cycle as a perturbation from the fixed point 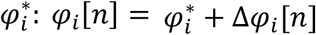, where *i* denotes the oscillator. We substitute these expressions and linearize the PRC around the fixed point at the locking phase using 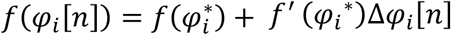. After eliminating all steady state terms, we obtain a system of three equations with four terms: Δφ_1_[*n*], Δφ_*N*−1_[*n*], Δφ_1_[*n*+1] to and Δφ_*N*−1_[*n*+1]. We then eliminate Δφ_*N*−1_[*n*] and Δφ_*N*−1_[*n*+1] to obtain a one-dimensional discrete map:

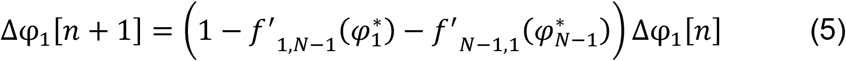

where *f*^′^_*i,j*_ are the slopes of the respective PRCs. The detailed derivation of this map is given in Appendix B. The map could also be defined based on Δφ_*N*−1_[*n*] and Δφ_*N*−1_[*n*+1] by eliminating Δφ_1_[*n*] and Δφ_1_[*n*+1].

For synchrony as a fixed point of this map to be stable, the perturbation must decay Δφ_1_[*n*+1] → 0 as n → ∞. Thus, the eigenvalue 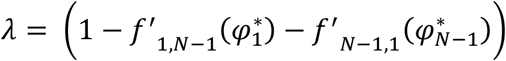 for the map in Eq. 5 must have an absolute value less than 1. We know the intrinsic period of a single oscillator and can obtain the period of the self-connected oscillator from the PRC of the single oscillator as 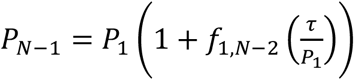. The PRC for the self-connected unperturbed network *f*_*N*−1,1_ is a function of the synaptic delay in the self-connection. Adding the fact that for synchrony the locking phase is *τ*/*P*_*i*_, we get the stability criterion described in the following theorem:

#### Theorem 1

**Stability of Synchrony in a Single Cluster**

*In a network of N identical neurons with synaptic delays and each receiving N-1 inputs, the following criterion is necessary for synchrony to be a stable solution:*

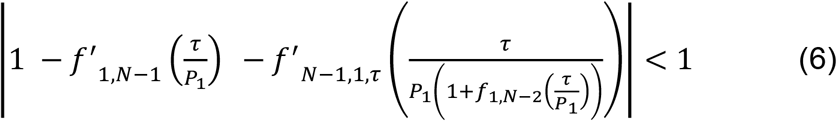

***Proof:*** Refer to Appendix B for the proof.

To test whether Eq. 6 made useful predictions in a system of pulse coupled oscillators in which all the assumptions in section 2.4 are honored, we used the emulator, or map with no predetermined firing order described in section 2.5. We modeled an all-to-all network of 25 identical model entorhinal cortical PV+ inhibitory neurons [14] described in Appendix A1. Optogenetic stimulation of these interneurons recruits interneuronal network gamma [36] with excitation blocked, thus these neurons can synchronize due to their connections with each other [37]. For illustrative purposes, we examined networks with two different values of the reversal potential of the inhibitory synapses since these two values are within the range of observed values for GABA_A_ synapses [38] but have very different synchronization properties [14,39].

The full set of ODEs for each oscillator was used to generate the PRCs in Fig. 3AB for each oscillator in response to a perturbation by the other oscillator (Fig. 2B). Fig. 3A sets E_GABA_ = -75 mV, which is called hyperpolarizing inhibition, and Fig 3B sets E_GABA_ = -55 mV, which is called shunting because the major effect is a conductance change with little effect on the membrane potential. The curves in blue represent the PRC of the single perturbed neuron due to the self-connected cluster *f*_*1,N-1*_ *(φ)* representing the rest of the network. The shapes for E_GABA_ = -75 mV and -55 mV are quite distinct. Since the PRCs for the self-connected oscillator are parameterized by the delay, the sets of points depicted in red color correspond to three points sampled around the locking phase (equal to the normalized feedback delay) in order to calculate the slope of the PRC *f*_*N-1,1,τ*_ *(φ)* for each delay *τ*. Once the PRCs were obtained, the network simulations were performed using only the PRCs and free-running period in the emulator hybrid phase model described in section 3.5.

**Fig. 3.**
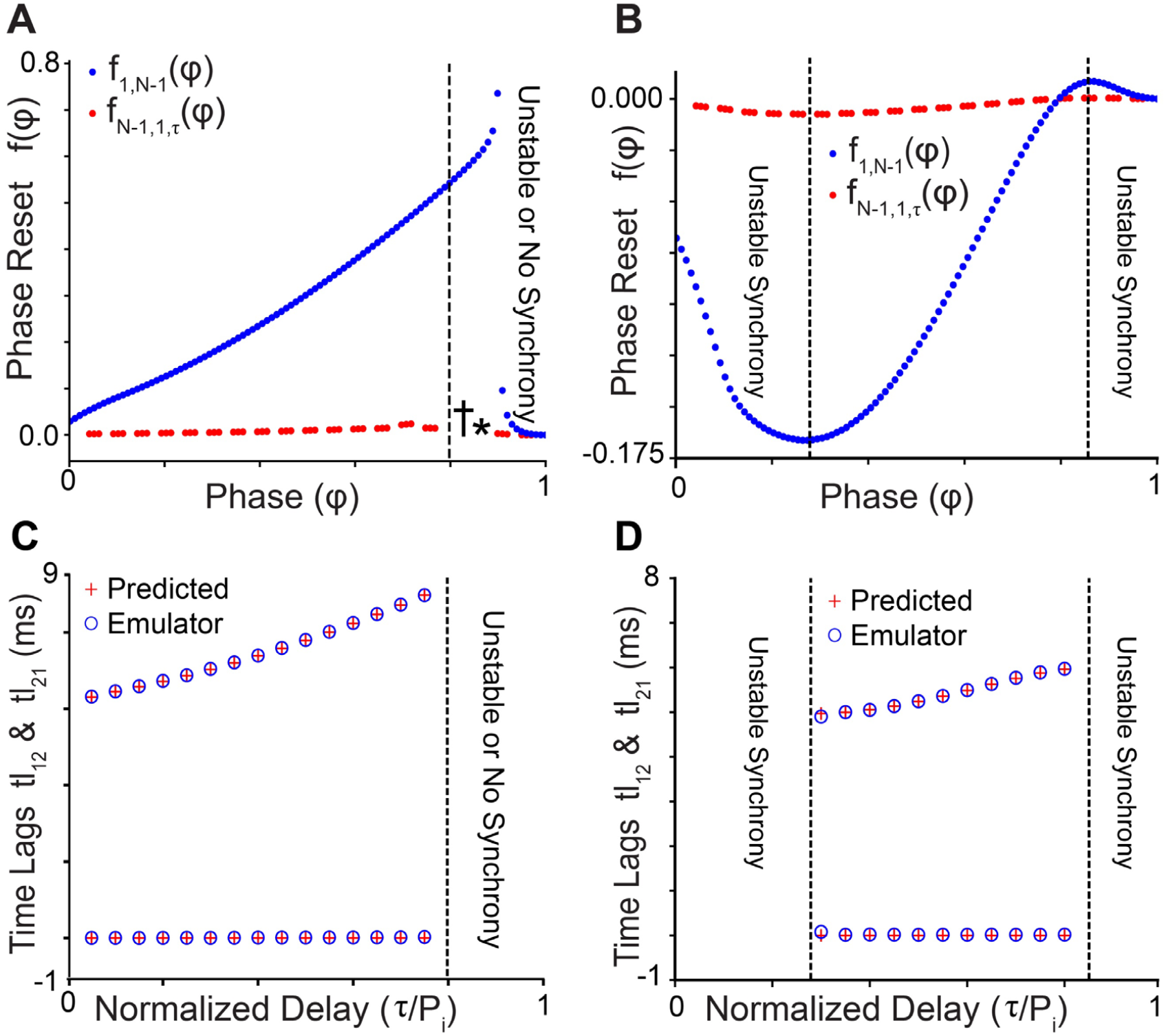
Predicted and Observed Periods for Global Synchrony. A. PRC of a single oscillator (blue) in response to a population spike in all other oscillators with E_GABA_ = -75 mV and of a self-connected oscillator representing the rest of the network. The set of points depicted in red color corresponds to three points from each PRC *f*_*N-1,1,τ*_ *(φ)* parameterized by the delay (equal to the locking phase) to give the slope at each delay *τ*. **\*** and **†** indicate the special cases with delays 4.8 ms and 5.1 ms, discussed in the case of existence of self-locking. B. Same as A with shunting reversal potential E_GABA_ = -55 mV. C and D. Predicted versus observed time lags in cases where synchrony was predicted to be stable using Eq.6. For synchrony, one time lag is zero (tl_12_) and the other (tl_21_) is the network period.

Fig. 3C and 3D show that the predictions of stability of synchrony are accurate because they agree with the emulator simulations. For synchrony, one time lag (tl_12_) is arbitrarily set to zero and the other (tl_21_) to the network period, calculated using: P^*^_N-1_ = (1 + *f*_*1*_*(φ\**)). To the right of the dashed line in Fig. 3A and outside the dashed lines in Fig. 3C, no time lags from the emulator map are presented because synchrony is predicted to be unstable. The eigenvalue of the discrete map calculated using Eq. 6 is greater than 1, because the delay caused a putative locking point to fall in the region of the PRC with negative slope, to the right of the discontinuity in Fig. 3A. Emulator simulations confirmed predictions that there was no convergence to synchrony in those cases. For the hyperpolarizing case presented in Fig. 3A and C, there were some discrepancies between theory and observations. For the cases with delays of 4.8 and 5.1 ms, denoted by a dagger and an asterisk respectively in Fig. 3A, the self-connected oscillator simulated in the ODEs could not phase lock itself. We explain these discrepancies in the next section.

### 4.2 Caveats on second order resetting: self-connected oscillator with delay

Despite our efforts to minimize second order resetting as described in section 2.1 and Appendix A, inputs applied at late phases often have effects on the subsequent cycle. In this section only, we allow second order resetting to be non-negligible. In order to incorporate second order resetting into our firing interval maps, we must assume that its effect is completed during the following stimulus interval (Fig. 1C) and does not bleed over into the next response interval. The result given in section 3.2 that a self-locking always occurs for short delays at *φ\**, where *τ* = *P*_*i*_*φ\** and *τ*< *P*_*i*_*(1+f’(φ*))*, ignores second order phase resetting. We observed some cases in which self-locking failed; therefore, we addressed the case of non-negligible second order resetting.

*Lemma*: For a 1:1 phase locked mode to exist, the stimulus interval at phase locking should be equal to the delay *τ* = *P*_*i*_*(φ* + f*_*2*_*(φ\**)), where *φ*\* is the phase at which an input arrives.

*Proof:* If there is no second order resetting, the existence of a stable 1:1 phase-locked mode is guaranteed at every delay less than the network period *τ* < P_i_(1+ *f*_*1*_*(φ*\*)), as shown in [23].

**Table 1.** Parameters for gating variables in Via model.

| | $m$ | $h$ | $n$ | $a$ |
| --- | --- | --- | --- | --- |
| $\Theta$ (mV) | -53.0 | -55.71 | 5.9 | 51.36 |
| $\sigma_1$ (mV) | 4 | -20 | 12 | 12 |
| $\sigma_2$ (mV) | -13 | 3.5 | -8.5 | -80 |
| $k_1$ (ms<sup>-1</sup>) | 0.25 | 0.012 | 1 | 1 |
| $k_2$ (ms<sup>-1</sup>) | 0.1 | 0.2 | 0.001 | 0.02 |

Figure 4A shows the firing interval map for a self-connected oscillator of the same type as in Figure 3A and C. With second order resetting, the steady-state stimulus interval ts[∞] becomes ts[∞] = *P*_*i*_*(φ* + f*_*2*_*(φ\**)), where *φ\** is the steady-state phase, and the input due to the self-connection arrives at a phase of τ/P_i_. For phase locking to exist, these two quantities must be equal to each other: *τ/P*_*i*_ = *P*_*i*_*(φ* + f*_*2*_*(φ\**)). Fig. 4B shows the predictions of the network period using the relation: P^*^_N-1_ = P_1_ (1 + *f*_*1*_*(φ\**) + *f*_*2*_*(φ\**)) compared to the observed results of the simulations of the ODEs of a self-connected oscillator. Figure 4C illustrates the graphical method to determine existence for *τ* = 5.1 ms such that *τ/P*_*i*_ ∼ 0.90. The green curve in the figure plots *P*_*i*_ *(φ* + f*_*2*_*(φ\**)) whereas the red dashed line has the constant value *τ/P*_*i*_ ∼ 0.9. The curves do not intersect because of the discontinuity in the PRC (at *φ* ≈ 0.9 in Fig. 1B), so no 1:1 phase-locking is predicted to exist at a delay of 5.1 ms, indicated by the asterisk in Fig. 4B and consistent with the ODE solutions of the self-connected oscillator shown in Fig. 4D. For other delays we find that the curves intersect (not shown here) predicting the existence of 1:1 phase locking. In addition to the existence criterion, the stability criterion for a self-locked oscillator with short delays considering second order resetting is *-1 < f*_*2*_*’(φ\**) <1 [7]. In all cases where existence was predicted, stability was also predicted.

**Fig. 4.**
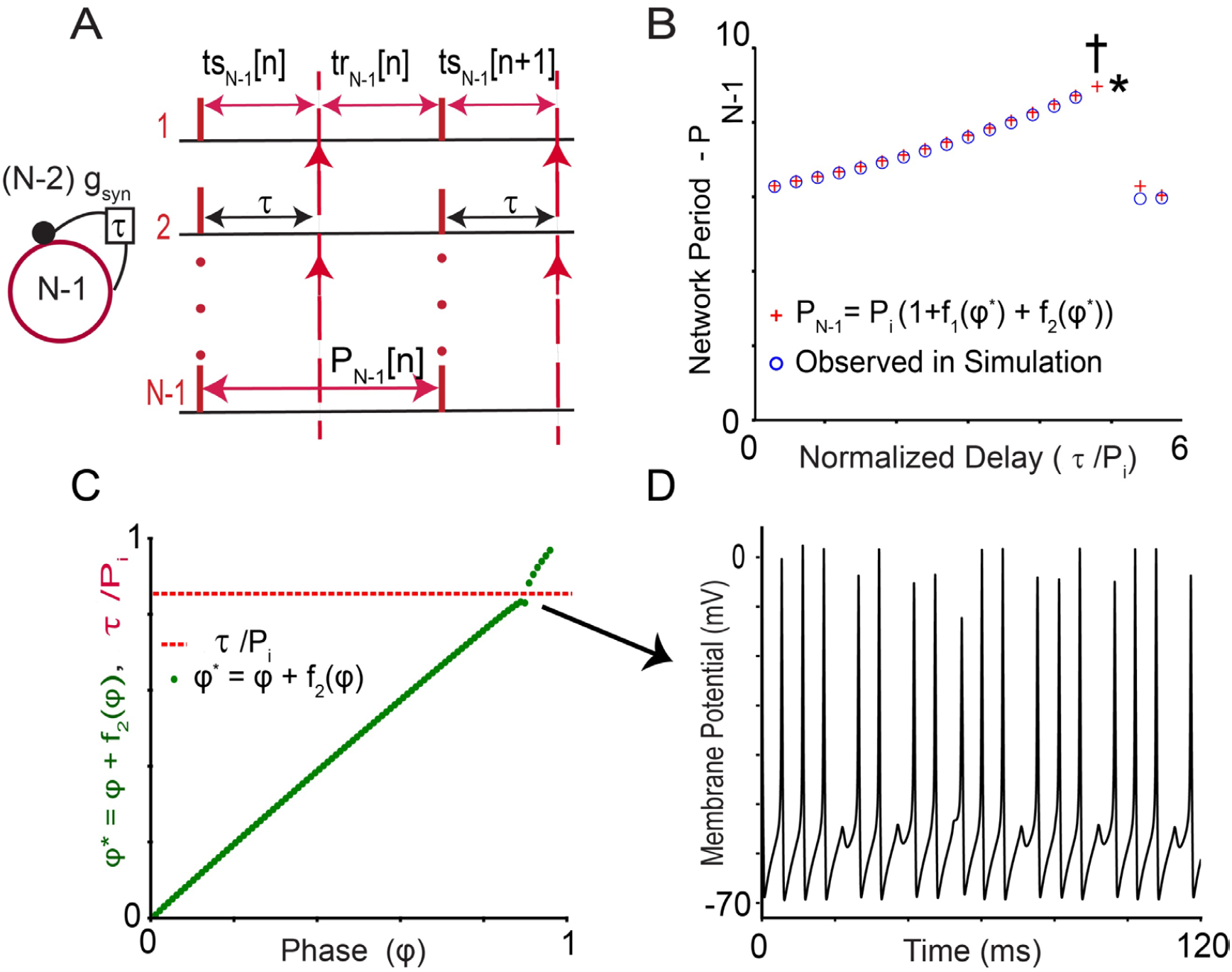
A. Interval map for the self-connected oscillator. The sub-cluster of N-1 neurons from a cluster of N neurons receive N-2 inputs after a delay *τ*. B. Plot of predicted and observed periods for various delays at which the inputs from self-connection arrive. The asterisk corresponds to the dashed red line in C for a delay of 5.1 ms. The dagger **†** corresponds to a delay of 4.8 ms. The assumptions are broken at that value as explained in the text, so an irregular mode like the one in panel D is observed instead of the predicted 1:1 locking. C. Steady-state stimulus interval ts[∞] = *φ* + f*_*2*_*(φ\**) plotted versus the phase *φ* does not intersect with the normalized delay of 5.1 ms, thus existence of self-locking is not predicted. D. Membrane potential trace during irregular firing mode with *τ* = 5.1 ms.

For a delay of 4.8 ms, existence of 1:1 self-locking is predicted by the intersection of curves; however, the ODE simulations produced an irregular self-locking similar to the one in Fig. 4D. This is the lone prediction failure, likely because the second order resetting due to one input may not have been completed before the next input was received, rendering the prediction indicated by **†** in figure 4B invalid. In a network of identical identically connected neurons, synchrony always exists due to the symmetry [40]. Fig. 4D shows that this synchrony need not be periodic.

### 4.3 Extension of the results for perturbation of more than one neuron

Eq. 6 is a necessary condition for stability, but it is not sufficient. For stability, synchrony must be robust to every perturbation in a small neighborhood of the fixed point of the map of the firing intervals. Additional perturbations to the synchronous firing pattern displace M neurons from the synchronous N-neuron cluster; each cluster is represented by a self-connected oscillator as in Figure 5A. The unperturbed cluster contains N-M neurons with N-M-1 self-connections, whereas the perturbed cluster with M neurons receives M-1 inputs due to self-connections. Extending Eq. 6 to the perturbation for M neurons (1 ≤ *M* ≤ *N*), we get a generalized criterion:

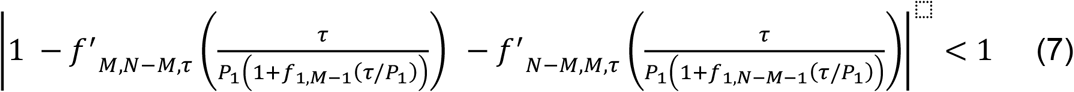

The PRCs of both oscillators are functions of the delay. To test the assumption that the single neuron perturbation from the network is the most destabilizing (Canavier and Achuthan, 2010), we used as an example the specific case from the model in the previous section which had the largest eigenvalue for the single neuron perturbation case, namely shunting inhibition with a delay of 0.3 ms. We then varied the number (M) of neurons perturbed from the 25-neuron network from 2 to 12 and used Eq. 7 to obtain the eigenvalue of the discrete map to predict stability. Figure 5B shows the slopes and eigenvalues for each case. As the M cluster gets larger and its input gets smaller, the slope of the PRC decreases. The slope for the partner N-M cluster increases because the cluster is decreasing in size and getting a larger input. For perturbations of 1 to 12 neurons, these increases and decreases offset each other rendering the eigenvalues quite consistent. They all are greater than 1 and predict that synchrony is unstable for each perturbation. Thus, for this example, perturbing any number of neurons serves to predict the stability of synchrony of the network.

**Fig 5.**
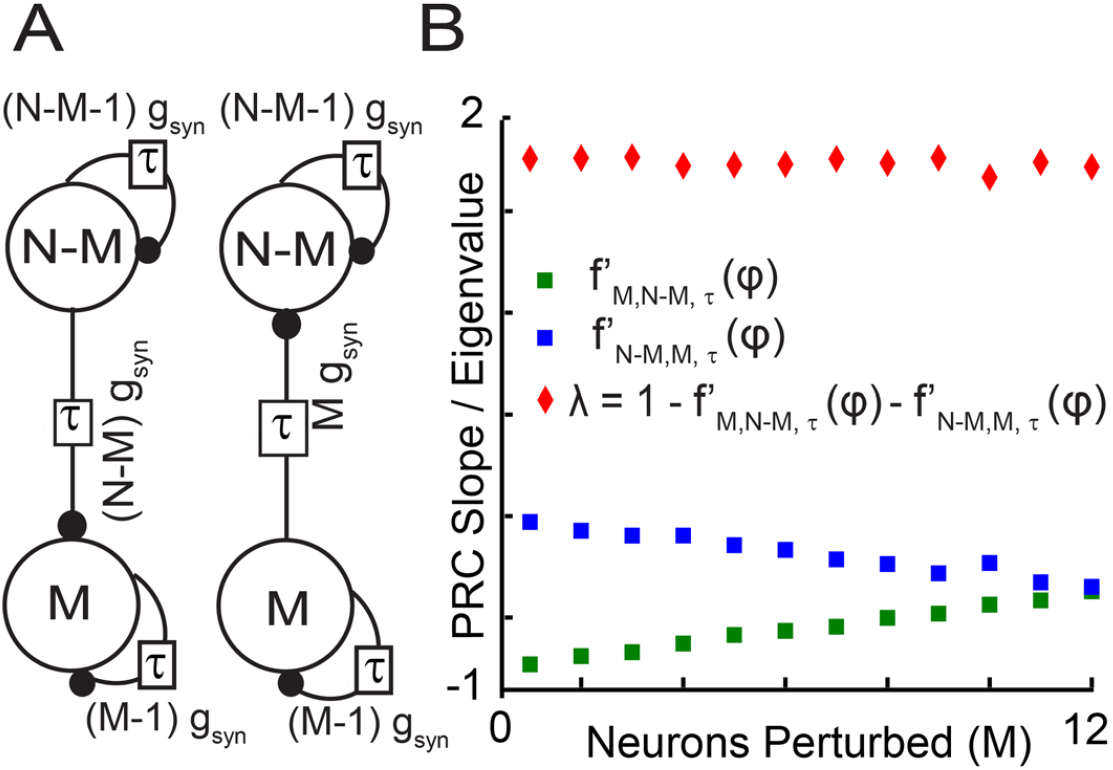
Multiple-Neuron Perturbations and Eigenvalues. **A**. Reduction of network to two oscillators with M neurons Perturbed form global synchrony. Both clusters are assumed to receive delayed synchronous feedback from themselves without autapses. **B**. For the example described in the text, the PRC slopes increase (decrease) as postsynaptic cluster size deceases (increases). These effects offset each other and the eigenvalue remains approximately constant.

### 4.4 Extension of the results for alternating firing of two synchronous clusters to include delays

In addition to global synchrony, two cluster solutions are often observed in homogenous systems of oscillators [33,34,41–44]. We assume once again that the stability of synchrony can be predicted by the response to the perturbation of a single neuron from a fully synchronized cluster, giving rise to the firing pattern illustrated in Fig. 6. The analysis can easily be generalized to perturbation of M neurons as in the previous section. If the clusters are not identical, the analysis should be repeated for a perturbation from cluster B. A two-cluster mode does not always exist because the between cluster terms must satisfy the existence criteria given in section 3.4 and determined by the firing intervals in Fig. 6 at steady state (unperturbed clusters).

**Fig 6.**
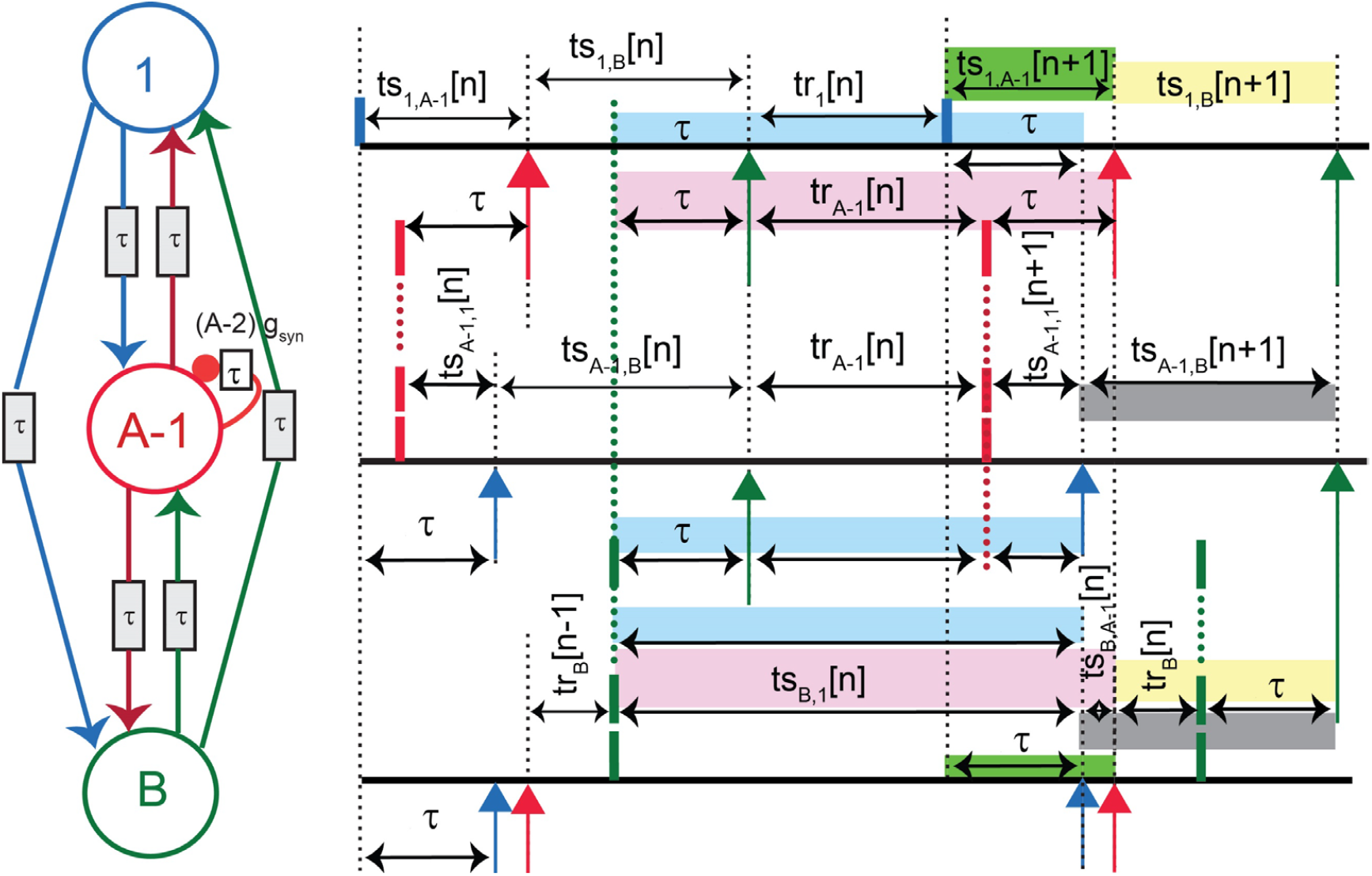
Firing intervals for perturbation of Two Cluster Network with Conduction Delays. Schematic firing pattern in which a single neuron is perturbed from one of the two clusters. These intervals are used to derive the discrete map through which the stability criteria of the two-cluster solution are established.

Recent work has shown that in networks with conduction delays composed of identical inhibitory neurons with the PRC shaped as in Fig. 1B, the presence of the discontinuity guarantees that if the solution of two identical clusters in antiphase does not exist, then all modes with two clusters of unequal size [34] and cluster modes with more than two clusters [45] do not exist.

To generalize previous results on pulse-coupled oscillators to the case in which two clusters synchronize in the presence of conduction delays, we consider both within cluster and between cluster terms. Within-cluster interactions in the presence of short delays take the form |*1 – f’(*τ*/P*_*i*_*) – f’(*τ*/P*_*i*_)| *< 1* as in sections 3.3 and 4.1. Between-cluster interactions take the form |*(1 – f’*_*A,B*_*(φ*_*A*_*\*)) (1 – f’*_*B,A*_*(φ*_*B*_*\*))*| *< 1* as in section 3.4 for two coupled oscillators. Previous studies that did not consider conduction delays were able to account for the observation [33] that an alternating firing pattern between two clusters can enforce synchrony within clusters that are not capable of synchronizing themselves in isolation [25,46]. The key insight is the multiplicative nature of the factors that scale the perturbation in separate intervals. The presence of three oscillators requires a two-dimensional discrete map with two eigenvalues. For two clusters without delay, the between-cluster term becomes |*(1 – f’*_*A,B*_*(φ*^*\**^_*A*_*)) (1 – f’*_*B,A-1*_*(φ*^*\**^_*B*_*))(1 – f’*_*B,1*_*(φ*^*\**^_*B*_))| *< 1* due to the perturbation in cluster A. We will show that this result is unchanged by delays except to parameterize the discrete system and the PRCs by the delay as needed. For no delays, the term containing the within-cluster effects for cluster A is scaled by the between-cluster term due to cluster B: *(1 – f’*_*A,B*_*(φ*^*\**^_*A*_). By analogy with [25], we expect the within-cluster term for the case with conduction delays to be |*(1 – f’*_*A,B*_*(φ*^*\**^_*A*_*)(1 – f’*_*A-1,1*_*(φ*_*A-1*_*\*) – f’*_*1,A-1*_*(φ*^*\**^_*1*_))| *< 1, where φ*^*\**^_*1*_*=*τ*/P*_*1*_ and *φ*^*\**^_*A-1*_*=τ/P*_*A-1*._ We prove these results below.

The sets of intervals shaded in each color (green, gray, yellow, blue and pink) in Figure 6 have equal length resulting in the following six equations:

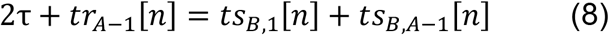

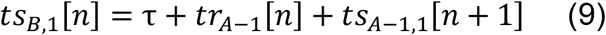

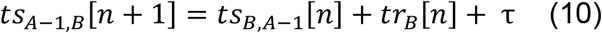

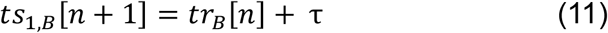

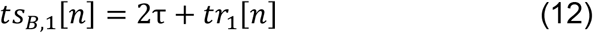

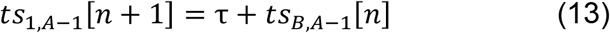

The key idea is that the perturbation between the A-1 cluster and the cluster of 1 must die out. This means the difference *ts*_1,*A*−1_[*n*]−*ts*_*A*−1,1_[*n*] must go to zero as n goes to infinity and both quantities approach *τ*. For stability, the deviations Δφ_*i,jj*_[*n*] from the steady locking phases 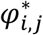 must also go to zero as n goes to infinity. At steady-state, the A-1 and 1 clusters merge into a single cluster with delayed feedback, rendering the network periods of all neurons equal. For within-cluster synchrony, the steady phase-locked network period of the single neuron is 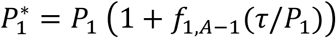 and the phase at which an input is received is 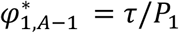. The free-running period of the self-connected A-1 cluster is *P*_*A*−1_ = *P*_1_ 1 +*f*_1,*A*−2_(*τ*/*P*_1_). The steady phase-locked network period of the A-1 cluster is 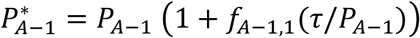, and the phase at which an input is received is 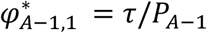. At convergence, cluster B receives a single merged input from A-1 and 1 so the steady state phases are: 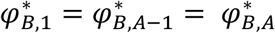. With these as steady states, we assume a perturbation about these and obtain the map for the phases at which an input is received in terms of phases in the previous cycle. The full derivation of the 2-dimensional discrete map and its eigenvalues is given in Appendix C, using the same techniques as in the previous section. The resultant 2-dimensional map is:

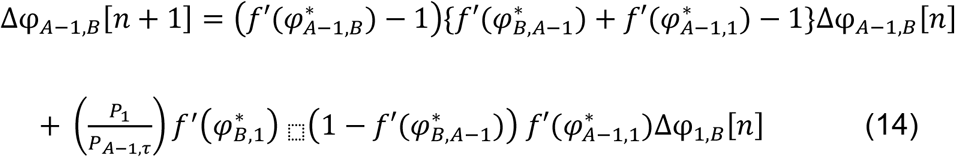

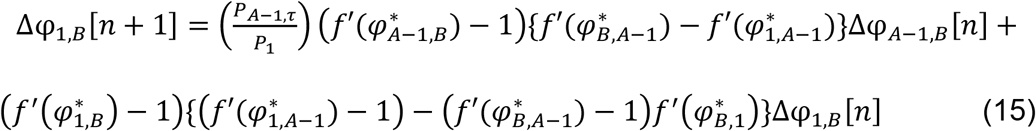

Eq. 14 and 15 constitute a discrete time 2-dimensional linear system of the form Δ[*n*+1] = *S*Δ[*n*], where S can be expressed as 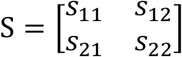. The terms *s*_11_, *s*_12_,*S*_21_ and *s*_22_ are defined as

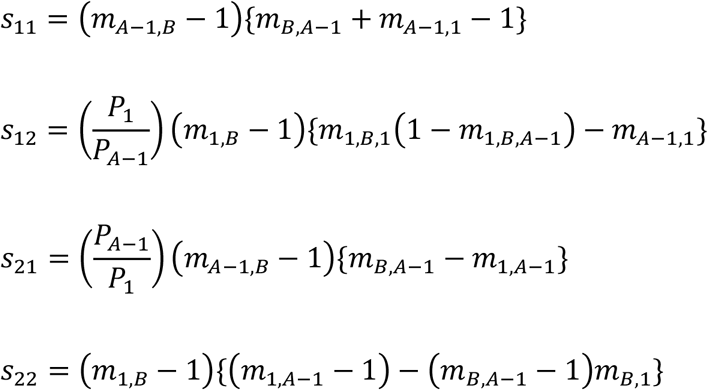

Based on this, the characteristic polynomial for the matrix S is given by

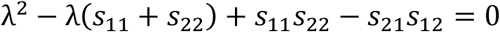

and the eigenvalues can be determined by finding the roots of the characteristic quadratic polynomial associated with this system. In contrast to the case without delays [25], the intrinsic periods of the oscillators appear in the discrete maps given in Eqs. 14 and 15. However, those two terms are multiplied and cancel each other out in the characteristic equation, so they do not contribute to the eigenvalues. For two self-connected clusters, the analysis must be repeated for a perturbation in cluster B. For the simplification of a cluster B with no self-connections, the criterion given by Eq. 14 and 15 suffices.

In order to simplify the 2-D map and gain intuition into how the eigenvalues are determined, we make the assumption that the PRCs of the subclusters on A to an input from B can be approximated by the PRC of the intact subcluster such that *f*_*A*−1,*B*_(φ_*A*−1,*BB*_) = *f*_1,*B*_(φ_1,*BB*_) = *f*_*A,B*_(φ_*A,B*_). In the specific example that we chose to illustrate the synchronization of one cluster by another, the green PRC for cluster A in Fig. 7B2 is very similar to the black dotted and dashed PRCs for the subcluster, so this assumption is justified.

**Figure 7.**
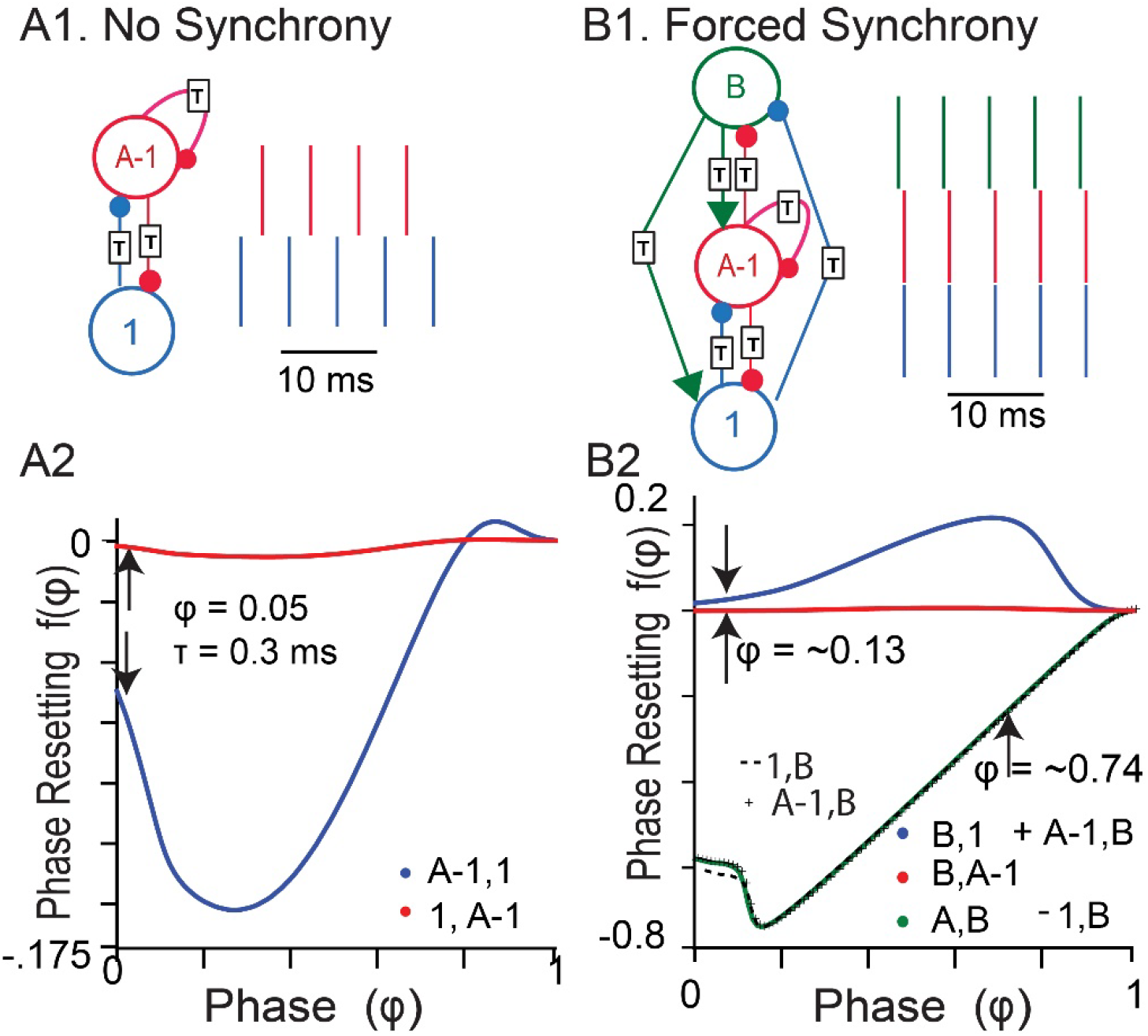
Two cluster synchrony A. A self-connected cluster of A neurons (red) cannot synchronize itself when one neuron (blue) is perturbed because the slopes of the PRCs at the locking point are negative. **B**. Mutual coupling to a population of excitatory cells (green) enforces synchrony.

Under this assumption, the eigenvalues are given in Eq. 18 and 19, and their absolute values must be less than one for stability.

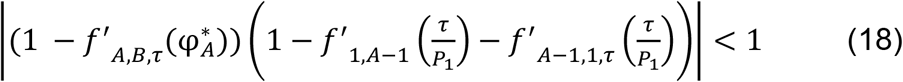

In Eq. 18, the large parentheses enclose the within-cluster synchrony term for cluster A which is scaled by the term representing forcing of cluster A by cluster B: 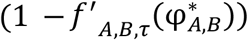. Since the slope of the green PRC in Fig. 7B2 is nearly 1 at the locking phase φ*, the absolute value of the eigenvalue becomes small even in the within cluster term within the large parenthesis has an absolute value greater than one. The other eigenvalue contains only between cluster terms:

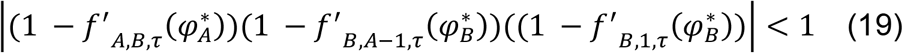

In contrast to the case without delays, the criteria in Eq. 18 and 19 suffice for perturbation of 1 from A independent of the order of firing.

As an example circuit in which to test the predictions of stability of synchrony for the two-cluster case with delays, we were motivated by experimental results showing that the parvalbumin positive fast spiking interneurons in the medial entorhinal cortex (mEC) cannot synchronize themselves with excitatory neurotransmission blocked, but synchronize at gamma frequency when reciprocal coupling to the excitatory population of neurons was intact [47]. We chose parameters for the mEC parvalbumin positive inhibitory model neurons for which global synchrony case was unstable as cluster A and a generic model of excitatory cortical neurons, the RTM model described in Appendix-A2 as cluster B. For simplicity, in this example we ignored self-connections among the excitatory population, consistent with experimental data on excitatory neurons in layer 2/3 entorhinal cortex [48].

The inhibitory neurons were connected by synapses that reversed at -55 mV with a delay of 0.3ms, which corresponds to the most destabilizing case for the cluster of inhibitory neurons as in the previous section. Figure 7A1 uses the PRCs of the interneurons (Figure 7A2) in the emulator to confirm prediction that they do not synchronize when a single neuron is perturbed from the inhibitory cluster. The arrows in Figure 7A2 show the negative destabilizing slopes of the PRCs at a delay of 0.3 ms (φ*=0.05). Figure 7B1 shows that adding reciprocal connections with the cluster of excitatory neurons in the emulator, using the PRCs in Fig. 7B2, results in synchrony between the single inhibitory neuron and the larger cluster of inhibitory neurons. Moreover, as stated above, Fig. 7B2 shows that the PRC *f*_*A*−1,*B*_(φ_*A*−1,*B*_) is indistinguishable from 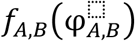 and 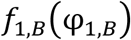 is barely distinguishable (and only in a small region of negative slope), thus justifying the approximation above that simplified the 2D discrete map whose eigenvalues are given in Eq 18 and 19. The excitatory neuron receives inhibitory input at a phase of 0.13 where the slope is small but positive, and the phase (φ*=0.74) at which the excitatory input reaches the inhibitory neurons has a strongly stabilizing slope ≈ 1. Thus, the between cluster scaling term *(1 – f’*_*A,B*_*(φ*_*A*_*\*)*) in Eq. 18 is nearly zero, bringing the unstable within-cluster eigenvalue of 1.75 down into the stable range with an absolute value less than one. Reciprocal coupling can be a powerful synchronizing influence.

## 5. Discussion

### 5.1 Generality

The results presented herein generalize to any system of pulse-coupled oscillators with delays, including synchronization of fireflies [4], cardiac pacemaker cells [49–51], circadian rhythms [52] and the engineering application of pulse-coupled sensor networks [53–56], provided the assumptions of pulsatile coupling are met.

### 5.2 Role of Delays in Neuronal Networks

In the nervous system, there is always a delay between an action potential in the pre-synaptic neuron and the postsynaptic potential observed at chemical synapses, due to the time required for an action potential to propagate from the axon initial segment to the axonal terminal and for the neurotransmitter to diffuse across the synaptic cleft and bind to the post-synaptic ionotropic receptors [57]. Delays in synchronization of inhibitory networks have previously been shown to promote synchrony [34,58]. The innovation herein is to use delayed self-connections to represent a perturbation of the network into smaller clusters to rigorously determine the stability of one and two-cluster synchrony. Delay-coupled neural networks can transition between synchrony and asynchrony as the level of excitation to the network is varied [59], a property that is not captured by systems without delay. The level of excitation controls the free-running periods of the neurons. Since the phase of the self-locking point occurs at the conduction delay normalized by the free-running period, the level of excitation also controls the slope of the PRC at the locking point and thereby controls stability. Moreover, the possibility that conduction delays themselves are plastic and contribute to synchronization and learning has been suggested to occur through regulation of axonal myelination [60,61].

### 5.3 Discontinuity in the PRC

Discontinuous PRCs have been previously observed [62,63]. Although the dynamical system representing the oscillator is continuous, the use of events such as the action potential to define the windows within which phase resetting occurs allows for discontinuities in the PRC. For example, the discontinuity at late phases in Fig. 1B occurs at the phase at which the perturbation cannot hyperpolarize the neuron sufficiently to stop the action potential from occurring, and the bulk of its effect occurs in the next oscillation cycle, resulting in second order resetting instead. The curves for both first and second order resetting are discontinuous. This discontinuity accounts for the non-existence of phase-locking of self-connected oscillator when the inputs arrive after a delay of 5.1 ms. As a result, the slope of the PRC becomes undefined at those points; therefore, phase locking does not exist at phases that fall in the discontinuity. However, it is clear that this discontinuity is highly repelling [34,45].

### 5.4 Higher-Order Phase Resetting

In the criterion for existence of phase-locking in a self-connected oscillator, we included second order resetting into the stimulus interval of the first cycle following the perturbation [33]. Otherwise, we ignored the second-order resetting, or any other higher order resetting. We showed that second order resetting can prevent the existence of self-locked synchrony, which is a prerequisite for the application of the stability criteria we present and limits their application when inputs are received too close to the time at which the next action potential would have occurred in the absence of a perturbation. This is not a serious limitation if one considers only delays that are much shorter than the network period. We reduced the decay time of the biexponential synaptic coupling term here to focus on the accuracy of the stability criteria. However, in the same inhibitory network as the one studied here, PRCs provide strong qualitative predictions of synchronization tendencies when longer, physiologically measured delays are incorporated into the network [14].

We do not incorporate third and higher order resetting into our simple pulsatile coupling framework, which is a limitation of our methods. Others [64] have considered the case in which an oscillator does not return to its original limit cycle by the time the next input is received. Some progress has been made on mean field methods for pulse coupled oscillators which could transcend these limitations [65] and render the results presented herein valid even in the presence of higher order resetting.

### 5.5 Perturbation of a single neuron from a cluster versus multiple neurons

From the empirical results in Fig. 5 for global synchrony in a network of inhibitory interneurons, we saw that the eigenvalues of the discrete map did not depend strongly on the number of neurons perturbed from global synchrony in that example. To guarantee stability, synchrony must be robust to all possible perturbations in some small neighborhood around the fixed point of the map. Therefore, if any specific perturbation is predicted to grow rather than decay, synchrony is unstable. However, the simplest perturbation (Eq. 6), that of a single neuron from a stable cluster, may provide useful insights regarding whether a system can synchronize, although technically Eq. 7 must also be considered.

### 5.6 Two-Cluster Applications to Neural Oscillations

There are numerous examples in the literature about how populations of excitatory neurons can synchronize themselves via mutual coupling with populations of inhibitory neurons [66–69] in a type of brain rhythm called pyramidal interneuronal network gamma (PING). Many of these mechanisms assume that the excitatory neurons are driven above threshold (mean driven) but inhibitory neurons are quiescent in the absence of excitation via their local excitatory neuron partners [70], whereas other assume that both populations are in the fluctuation driven regime below threshold [71–73]. We propose a third mechanism by which the excitatory cells can synchronize tonically firing interneurons that cannot synchronize themselves, as in Fig. 7. During theta oscillations, the firing rate of fast spiking interneuron increases [74], thus it is possible that interneurons are in the mean driven regime during theta nested gamma. In optogenetically-driven theta-nested gamma in a slice preparation [47], the interneurons are indeed tonically active during the peak of theta with excitation blocked. In that study, the interneurons only synchronized with excitation intact. Under similar optogenetic drive, we have observed some gamma synchrony in the absence of excitation; however, recruiting excitatory cells into optogenetically-driven theta nested gamma [37] increases gamma power, perhaps by a similar mechanism as in Fig. 7.

Another application of these results is to the case in which a single population can no longer support global synchrony but breaks into two clusters, doubling the observed population frequency. For example, the beta peak observed in slices of entorhinal cortex [75] has been suggested to be generated when a synchronized population of entorhinal stellate cells breaks up into two clusters firing at theta frequency in antiphase with each other [42]. Similarly, the frequency of sleep spindles at twice that of the delta rhythm has been hypothesized to result from the population splitting into two clusters firing at delta frequency in antiphase with each other [43]. Finally, based on the observation that individual cells in hippocampal area CA3 can fire no faster than 300–400Hz, fast ripple activity at 600 Hz was suggested to result from clusters in antiphase [44].

## 6. Conclusions

We extended the criteria for stability of one and two cluster synchrony in networks of pulse-coupled oscillators to include conduction delays shorter than the intrinsic period using self-connected oscillators to represent perturbed clusters of oscillators. Just as in the case of two oscillators, adding delays changed not only the locking point but also the criteria for stability of both global synchrony and of synchrony in two alternating clusters.

## Acknowledgements

This work was supported by the National Institutes of Health grant R01NS054281. The funding agency had no role in the collection, analysis and interpretation of data, in the writing of the report or in the decision to submit the article for publication.

## Conflict of Interest

The authors declare no conflict of interest.

## Appendix A: Conductance Based Neuron models for Simulations

### A.1. Via Model for Inhibitory Neurons

In this study, we examined a model of a representative fast spiking parvalbumin (PV+) inhibitory interneuron in layers 2/3 of the medial entorhinal cortex [14]. This model was calibrated based on data the passive and active properties of these neurons, which have Hodgkin’s class 2 excitability meaning they cannot sustain repetitive firing below a threshold frequency. The single compartment model neurons have five state variables: the membrane potential, *V*, and four gating variables (m, h, n, and a) which use the same kinetic equations as the original Hodgkin-Huxley model [76,77], but with different parameters tuned to replicate the dynamics of fast spiking neurons in the medial entorhinal cortex and given in Table 1 of [14]. We also included two delayed rectifier K_+_ currents (I_Kv1_ and I_Kv3_).

The differential equation for the membrane potential *V* is given by:

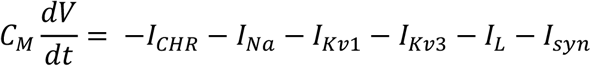

where *C*_*M*_ is the membrane capacitance, *I*_*CHR*_ is the simulated constant current replicating the optogenetic drive, *I*_*Na*_ is the fast sodium current, *I*_*L*_ is the leak current, *I*_*syn*_ is the GABA_A_ synaptic current. The ionic-current equations are:

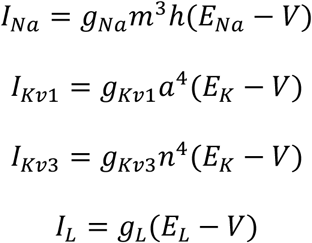

with *E*_*Na*_ = 50 mV, *E*_*K*_ = -90 mV and *E*_*L*_ = -65 mV. The dynamics of the gating variables are given by 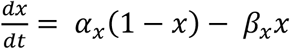 for the activation variables (m, n, a) and by 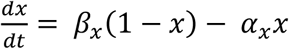 for the inactivation variable h, where *α*_*x*_ = *k*_1*x*_ (*θ*_*x*_ −*V*) / (exp((*θ*_*x*_ −*V*) / *σ*_1*x*_) −1) and β_*x*_ = *k*_2*x*_ exp(*V* / *σ* _2*x*_) with parameters in Table 1.

All the synaptic parameters except one (synaptic decay constant) were calibrated according to [14] with a biexponential waveform in conductance with time constants of *τ*_1_ = 0.3 and *τ*_2_ = 0.45 ms and a peak conductance of 1.65 nS: *g*(*t*) = *F*(exp(−(*t* − *τ*)/ *τ*_1_) − exp(−(*t* − *τ*)/*τ*_2_)), where F is a normalization factor that sets the peak to value 1. For the self-connected oscillator, this conduction waveform was initiated after a conduction delay *τ* by each spike in the neuron. The synaptic decay constant *τ*_1_ was 2.0 ms in [14] but was reduced to 0.45ms to reduce second order resetting. Two values for the synaptic reversal potential (*E*_*REV*_ = -75 mV and *E*_*REV*_ = -55 mV) were used to produce two very different phase resetting curves and the synaptic delay was varied to correspond to phases from 0.05 to 0.95 to test our stability criteria.

### A.2. RTM Model for Excitatory Stellate Cells

To simulate a population of excitatory neurons, we used the Reduced-Traub-Miles model (RTM model) of an excitatory neuron in the rat hippocampus [15] which is a slight modification of the model of Ermentrout and Kopell [78]. The single compartment model neurons have five state variables: the membrane potential, *V*, and four gating variables (m, h, and n) which use the same kinetic equations as the original Hodgkin-Huxley model with parameters given in Table 5.1 of [79]. Instead of applying a constant current to induce repetitive firing, we used a constant excitatory conductance to simulate channel rhodopsin drive as in the Via Model above.

## Appendix B: Derivation for Stability of Synchrony in an All-to-All Network

Expanding the intervals in Equations 2-4 using the definitions of the stimulus and response intervals of each oscillator, substituting the phases as perturbations from the steady state and linearizing the PRCs we get

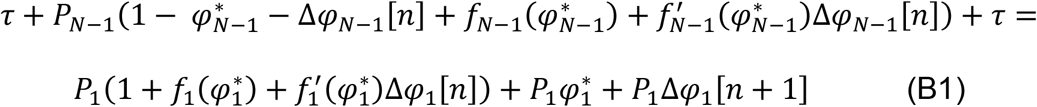

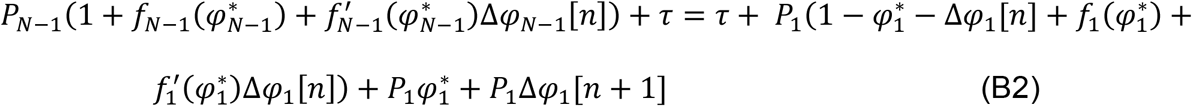

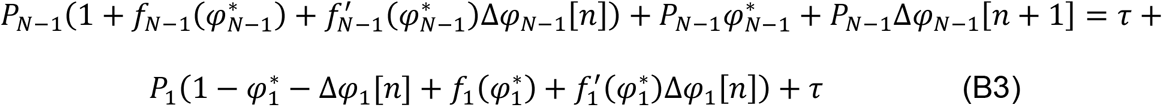

Denoting 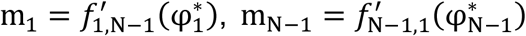, and cancelling the steady-state terms we get the equations

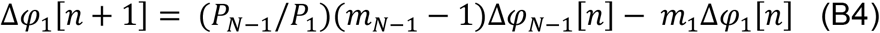

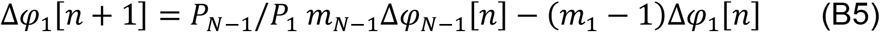

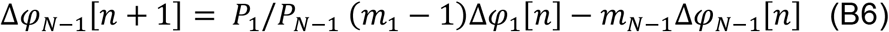

Now, substituting the expression for Δφ_*N*−1_[*N*] from equation (B5) into equation (B4), we get

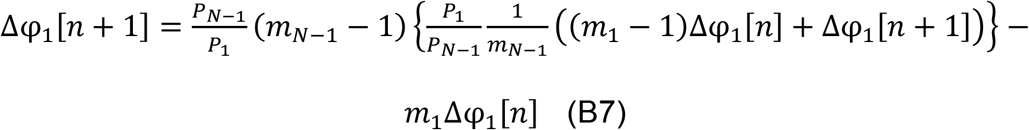

The periods get cancelled and multiplying each side by *m*_*N*−1_, re-arranging and simplifying, we get the one-dimensional discrete linear map for the perturbation of the single oscillator:

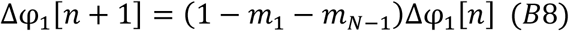

Note that if we used (B5) and (B6), we would have eliminated Δφ_1_[*n*] and the map would be in terms of Δφ_*N*−1_[*n*+1].

The steady locking phases in a synchronous network are

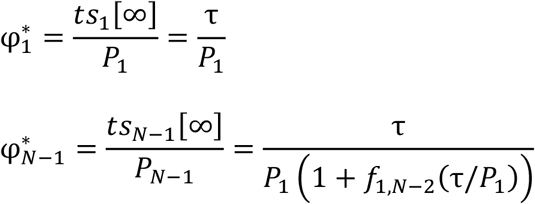

Substituting the values for the slopes *m*_*i*_ of the PRCs, the map at a synchronous state is given by

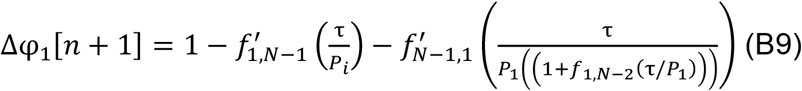

Since the PRC *f*_*N*−1,1_is measured by simulating the unperturbed cluster using self-connections and delay, the slope 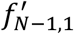 becomes a function of delay and thus the map becomes:

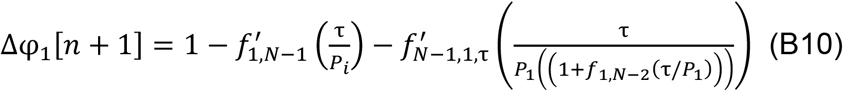

## Appendix C: Derivation for Stability of Clusters with Delays

We consider only delays that are small relative to the network period and equal to each other. Substituting the expressions for stimulus and response intervals into Eq. 8, we get

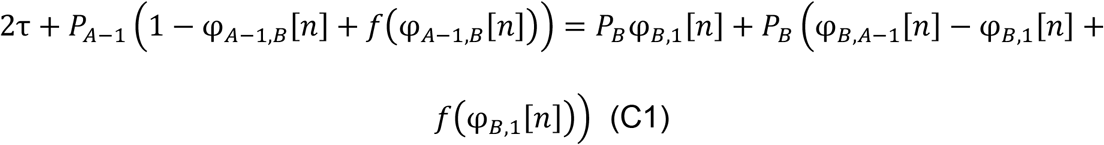

As discussed in the results section, the steady state here is the two-cluster solution, in which each cluster is composed of neurons that fire synchronously, and the two clusters fire in alternation. Substituting the perturbation from the steady-state phases, linearizing the PRCs about the fixed point, and canceling out the steady state terms on either side of the equation as in Appendix B, we get

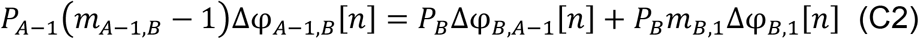

**Note:** As in Appendix B, we denote *m*_*i,j*_ = *f*′(φ_*i,j*_) for convenience.

Rearranging the above equation, we get:

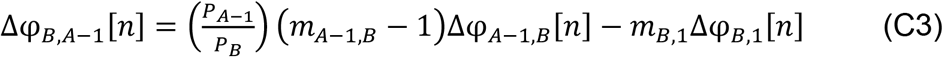

Writing Eq 12 in terms of the phases, we get

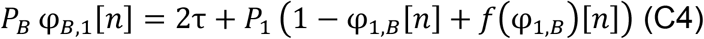

Rewriting the phase as a perturbed phase, linearizing about the fixed point and canceling out steady state terms on each side of the Eq C4, we get

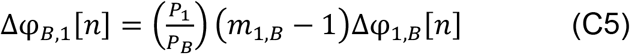

Substituting Δφ_*B*,1_[*n*] from equation (C5) in (C3), we get

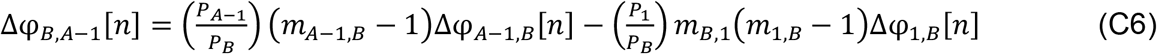

Substituting phases into Eq 9, we get

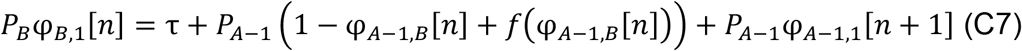

Rewriting the phase as a perturbed phase, linearizing about the fixed point and canceling out steady state terms on each side of the Eq C4, we get

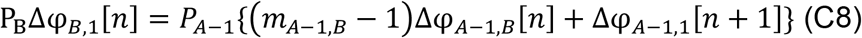

Rearranging the above expression, we get

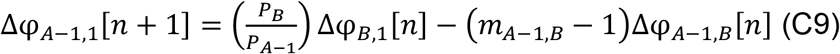

Substituting for φ_B,1_[*N*] from Eq C5 into Eq C9, we get

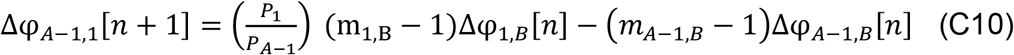

Substituting the definition of the intervals in Eq 13 terms of the phases, we get

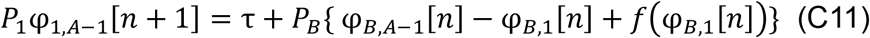

Rewriting the phase as a perturbed phase, linearizing about the fixed point and canceling out steady state terms on each side of Eq C4, we get

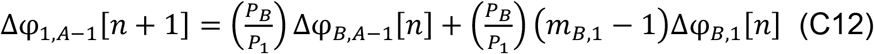

Substituting for Δφ_B,A−1_[*n*] in the above equation from (C6) and simplifying, we get

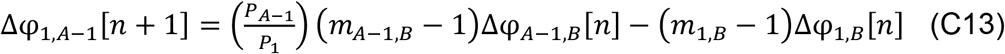

Substituting phases in Eq. 10, we get

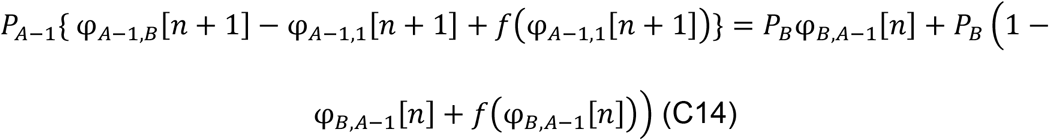

After Rewriting the phase as a perturbed phase, linearizing about the fixed point and canceling out steady state terms on each side, by rearranging the terms we get

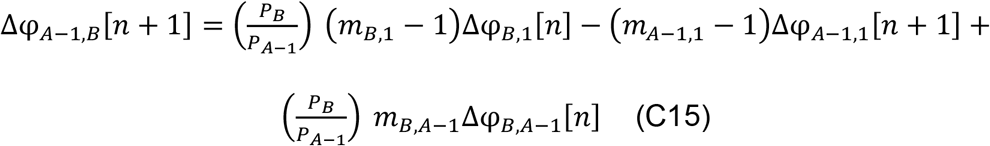

Substituting for Δφ_A−1,1_[*n* +1] in the above equation from equation (C10) and for Δφ_B,1_[*n*] in the above equation from (C5), we get

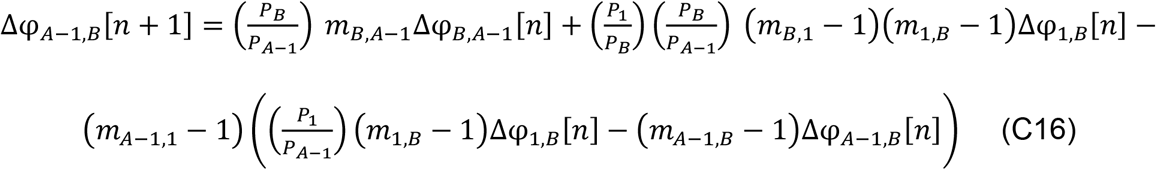

Rearranging and simplifying, we get the following:

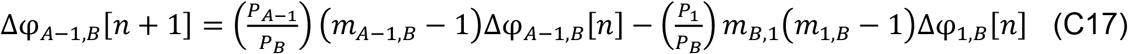

Now, we substitute for Δφ_B,A−1_[*n*] in the above equation from (C6), rearranging and simplifying results in the first equation of the map

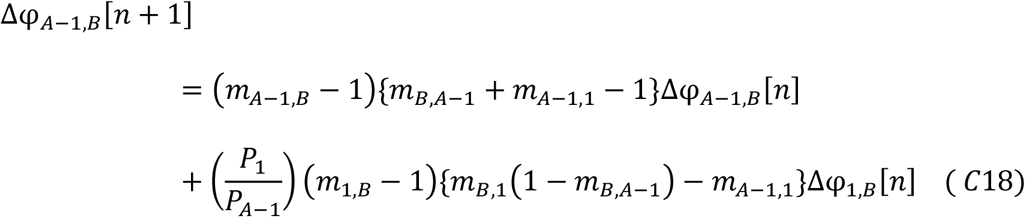

Substituting the phases for the intervals in Eq 11, linearizing the PRCs about the fixed points, cancelling the steady state terms and rearranging them, we get

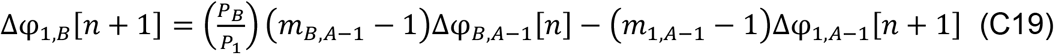

Substituting for φ_B,A−1_[*n*] from equation (C6) and for φ_A−1,1_[*n*] from equation (C10), we get

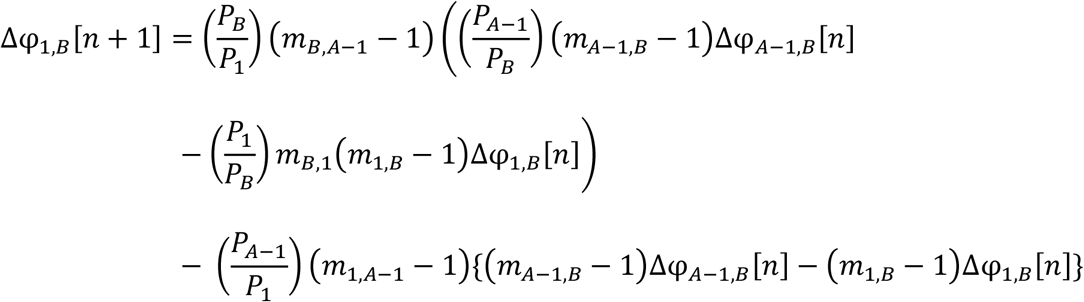

Cancelling the terms and rearranging, we finally get the second equation of the map

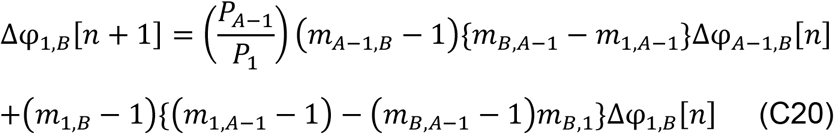

## Notes

### Competing Interest Statement

The authors have declared no competing interest.

### Summary of Updates

1. We have included some more literature concerning pulse coupled oscillators and PRCs. 2. We have made the derivations more brief. 3. Some notation that were used across the manuscript were made consistent.

